# Bypassing Current Biosecurity Screens with AI Designed Proteins and Closing the Gap with Edge-AI Functional Screening

**DOI:** 10.64898/2026.10.14.759233

**Authors:** Mackenzie Noon, Grace Oualline

**Author notes:** These authors contributed equally to this work.

## Abstract

DNA synthesis machines are vital tools in medicine and research, but they can be misused to produce sequences which are transcribed and translated to produce toxic proteins. Synthesis providers may screen orders for dangerous sequences, but most screening software performs sequence-based alignment, which is vulnerable to AI redesign attacks. Additionally, software-only synthesis screening requires the operator of the machine to be trusted. We present *Capiti*, an edge-deployed screening system that runs with no internet connection, listens directly to a synthesizer’s valve control signals, and aborts a run when it detects incipient pathogenic function. Rather than performing sequence alignment, *Capiti* is specifically trained to recognize fold. ProteinMPNN was used to generate putatively functional variants of known pathogenic proteins, and a CNN was trained to separate them from active-site knockouts, scrambles, and decoys. We release two models, *Capiti-E* (known pathogenic proteins) and *Capiti-C* (resistance genes for antibiotics of last resort), alongside emusynth, a DNA synthesizer emulator. On our test sets, IBBIS commec and SeqScreen incorrectly passed 18.5% and 44.7% of putatively functional pathogenic variants respectively; *Capiti* passed 0.24%.

**Graphical Abstract:** 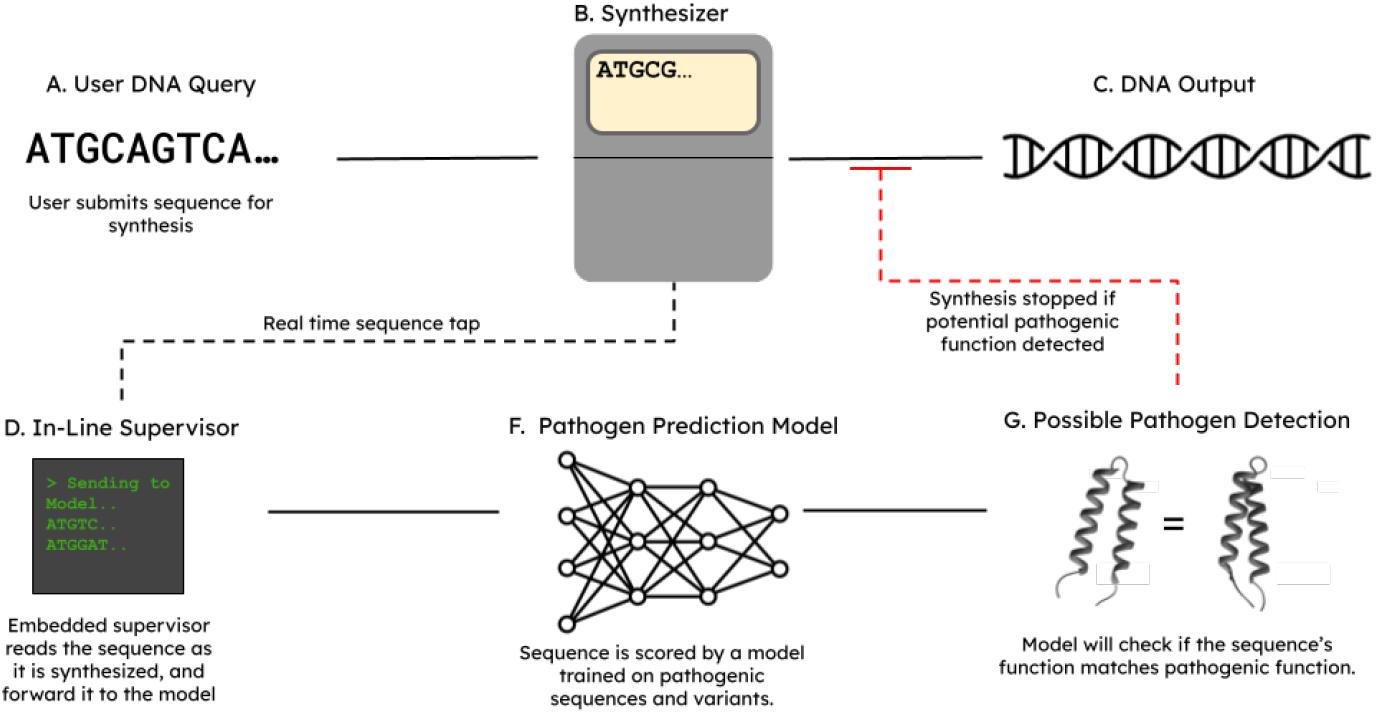

## Introduction

Benchtop DNA synthesizers have become essential tools for molecular research. Current enzymatic benchtop devices can synthesize up to 300 base pairs, and over the next 2–5 years, this is projected to reach 7,000 bp, about the size of the smallest viruses (Robert F. Service 2023). As synthesis length scales, so does misuse potential. Unlike traditional synthesis services, where screening is performed at a centralized location and risk management can be centralized (Laird et al. 2025b), the risk posed by benchtop synthesis machines is diffuse.

Several platforms have demonstrated the feasibility of biosecurity screening for benchtop synthesizers. SecureDNA and IBBIS commec use k-mer lookup and HMM-based profile search, respectively, to flag sequences of concern prior to synthesis (“SecureDNA Research,” n.d.); (Laird et al. 2025a). However, a cross-sector team demonstrated that AI-designed synthetic variants of known toxins evade homology-based screening (Wittmann et al. 2025).

In addition to challenges with diffuse risk and AI design, there is the issue of which sequences are included in biosecurity systems, and the notable absence of high-risk antimicrobial resistance (AMR) genes. AMR is a first-order public health crisis. Bacterial AMR was directly responsible for 1.27 million deaths in 2019 and contributed to 4.95 million deaths that year (“Antimicrobial Resistance” 2023). Forecasts from the GRAM Project suggest AMR will cause 39 million deaths between 2025 and 2050 (“Antibiotic Resistance Could Cause 39 Million Deaths between Now and 2050” 2024). The primary driver is the misuse and overuse of antimicrobials in humans, and animals, which accelerates the emergence of drug-resistant pathogens (“Antimicrobial Resistance” 2023). Antibiotics are given to livestock to prevent disease and improve growth margins, and this generates sustained selective pressure on bacterial populations and drives the emergence of new resistance genes. *Of particular concern are resistance genes to antibiotics of last resort*. If these genes become widespread in pathogenic bacteria, infections that are currently treatable will become untreatable by any available antibiotic class (“WHO Bacterial Priority Pathogens List, 2024: Bacterial Pathogens of Public Health Importance to Guide Research, Development and Strategies to Prevent and Control Antimicrobial Resistance” 2024). Agricultural operations seeking to protect livestock from resistant infections (Schoenmakers 2020), and adversarial actors looking to engineer pathogens, present realistic misuse scenarios for benchtop synthesis of these genes.

**Our main contributions are:**

1. *Capiti*: a hardware and software biosecurity system that intercepts DNA synthesis machine valve control signals, identifies incipient pathogenic function mid-synthesis, even when obscured by AI protein design tools, and halts synthesis before completion. Runs on a Raspberry Pi for edge deployment.
2. *Capiti-C* : a lightweight machine learning model for on-device antibiotic resistance screening, specifically targeting resistance genes to antibiotics of last resort, allowing legitimate use while preventing biorisk.
3. *Capiti-E* : a lightweight machine learning model for on-device identification of diverse pathogenic functions.
4. Emusynth: an Arduino-based emulator of a DNA synthesis machine liquid handler microcontroller, used for development and testing.
5. Discovery of vulnerabilities to AI redesign in existing nucleotide screening tools.

## Related Work

Benchtop sequence screening systems like SecureDNA, SeqScreen, and IBBIS commec could address the biosecurity risk posed by benchtop synthesis machines through their lookup-based approaches (“SecureDNA Research,” n.d.); (Balaji et al. 2022). Though groundbreaking, these solutions have several limitations. Variously:

### Issues with current pathogen screening systems

1. **Many require internet access**, which is problematic as it increases the attack surface and is incompatible with air-gapped or low-connectivity environments. Pre-issuing synthesis certificates is a partial solution but imposes a significant burden on the researcher. Remote server latency also introduces a timing problem: by the time a flagged sequence is identified, synthesis may already be underway or complete.
2. **Homology-based matching is susceptible to AI-driven redesign**. Proteins can be engineered to have minimal sequence similarity to known threats while fully preserving biological function, and these variants can bypass existing screening approaches (Wittmann et al. 2025).
3. **Existing approaches neglect antibiotic resistance**. Benchtop synthesizers provide meaningful uplift to actors seeking to engineer antibiotic-resistant pathogens, yet AMR genes are excluded from most screening databases and threat frameworks.

We present *Capiti*, an edge-ai biosecurity solution for benchtop DNA synthesis machines to address these challenges.

### How *Capiti* addresses current issues

1. *Capiti* **can run without internet access** on low-cost hardware, physically separate from the primary synthesis controller. Even in the event of a remote compromise, which is unlikely given the absence of network access, *Capiti* cannot direct synthesis, but can only interrupt it, providing a safety intervention without expanding the attack surface. This architecture also resolves the latency problem, as *Capiti* operates locally and identifies pathogenic sequences mid-synthesis in real time.
2. *Capiti* **matches pathogenic function rather than matching against a pre-defined library**. We used ProteinMPNN to redesign pathogenic sequences, and trained *Capiti* to catch them. This makes detection robust to AI-redesign attacks, since *Capiti* learns functional features which are preserved even when sequence identity is not.
3. *Capiti-C* **extends coverage to last-resort-antibiotic resistance genes**, a threat category largely absent from existing screening platforms.

*Capiti* addresses several threat models, but is most effective against a remote, persistent, advanced attacker seeking to synthesize sequences of concern on a compromised benchtop device, as defined in (Langenkamp 2024). In this scenario, an attacker gains remote access to a synthesis machine and attempts to produce a pathogenic sequence. *Capiti* intercepts the valve control signals mid-synthesis and aborts the run before completion. *Capiti* prevents spoofing by directly listening to valve signals, rather than relying on any external source of truth. The threat models addressable by *Capiti* could be substantially extended through the addition of tamper-proofing measures and cryptographic logging, which we discuss in future directions.

## Methods

### Construction of Hardware for Benchtop Synthesis Intercepter and Screening Platform

First, we constructed a test rig (shown in Figure 1 and schematized in Figure S1) consisting of an Arduino UNO running Emusynth and a Raspberry Pi running *Capiti*. The two devices communicate over a nucleotide bus of five wires, one for each of the four nucleotide valve signals and one for the Tetrazole activator, along with two additional control lines. An end-of-synthesis line pulses at the completion of a synthesis run, and an interrupt line pulses when *Capiti* makes a high-confidence pathogenic sequence call, aborting the run. An LCD screen displays Emusynth status and indicator LEDs were included for debugging.

**Figure 1:**
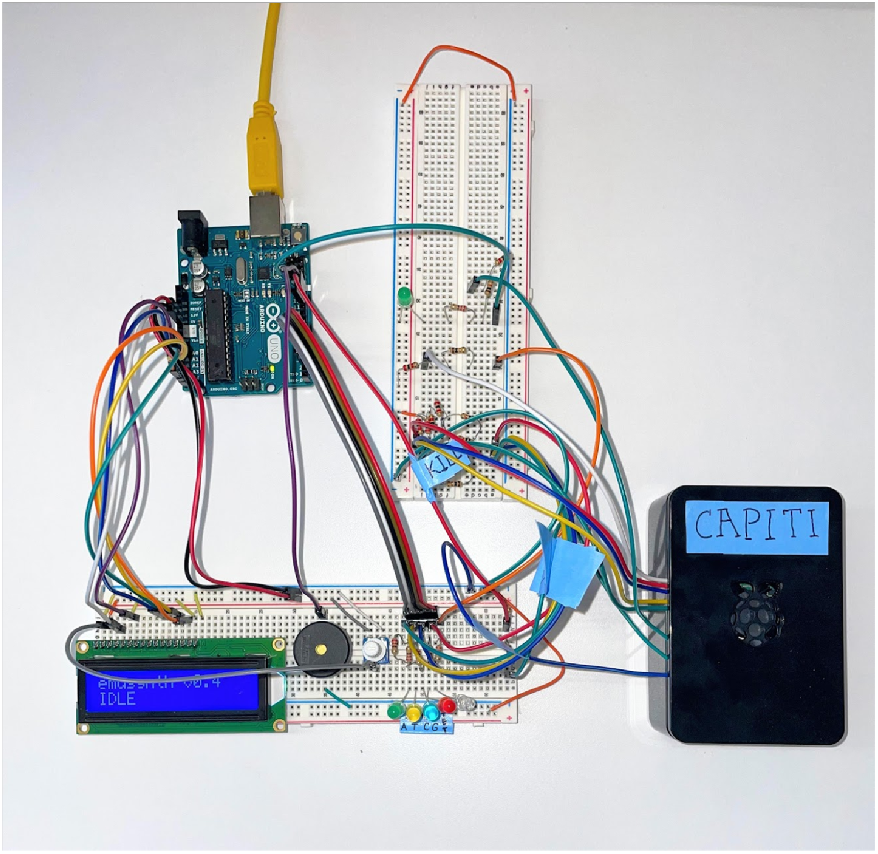
Assembled test rig of *Capiti* and *emusynth*.

Emusynth emulates the solenoid valve pulses of the Applied Biosystems 3400 DNA Synthesizer, selected for its widespread use and publicly accessible documentation (“Applied Biosystems 3400 DNA Synthesizer User Guide” 2010); (Pubcompare 2026). The approach generalizes in principle to most other synthesizer models. Emusynth consists of a firmware core running on the Arduino and ancillary Python scripts running on a laptop. The firmware accepts a minimal command set over a serial connection, `LOAD, RUN, STOP`, and `STATUS`, plus a `DIL` command to scale synthesis speed for testing purposes, and streams emulated pulses onto the nucleotide bus. Streaming was necessary because many of the sequences tested exceeded the Arduino’s onboard memory. Emusynth emulates all five synthesis steps: Detritylation, Coupling, Capping, Oxidation, and Washing, though only coupling step pulses are used downstream, as this step determines which nucleotide is added to the growing chain. *Click here to see a video demonstration of emusynth*.

### Creation of a database of pathogenic proteins that match function and not sequence

To train *Capiti*, we assembled datasets of proteins with known pathogenic function. Key functional residues were fixed and a library of putatively functional variants was generated using ProteinMPNN (Dauparas et al. 2022). This functioned as our positive set. We then created a “negative” set from six categories: two kinds of active site knockout strategies, alanine scanning (`ala_scan`) and combined knockout (`combined_ko`); two scrambling strategies, full scramble (scramble) and partial perturbation at 30% (`perturb30`); and unrelated sequences (`random_decoy`). We also trialed including hand-picked nonpathogenic sequences from the same protein family (`family_decoy`) as negatives. Sequence truncation was applied during training and test set generation to train the model to recognize pathogenic function from incomplete sequence fragments, reflecting realistic synthesis interception conditions.

**Fig 2:**
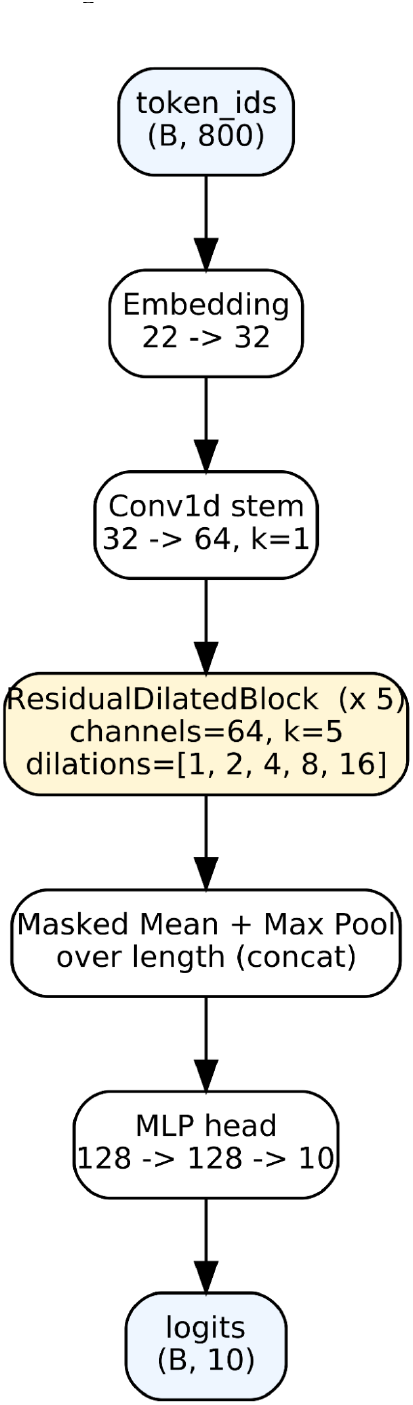
Capiti Architecture

**Fig 3:**
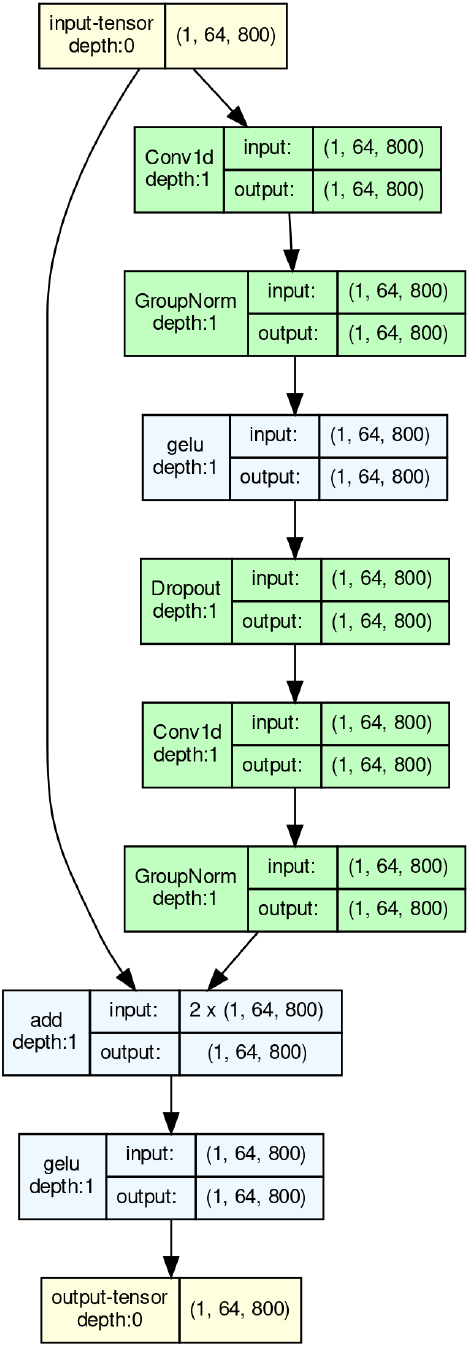
Residual Dilated Block detail

#### Next, we designed and trained several versions of *Capiti*

We designed *Capiti*’s network architecture to be sophisticated enough to learn pathogenic function, but light enough that inference can be run on embedded systems. Inspired by previous work that demonstrated excellent performance of simple Convolutional Neural Networks for sequence modeling tasks (Bai et al. 2018), and our desire to keep the model lightweight, we iterated on a simple CNN design. Dilation allowed the model to learn distant features without computationally expensive attention heads, and residual connections allowed the blocks to specialize and mitigated issues with vanishing gradients. ProteinMPNN variant generation and *Capiti* model training were performed on an NVIDIA RTX™ 5000 Ada Generation GPU.

We provide two pre-trained versions of *Capiti*, one for known pathogenic proteins, and one for resistance genes for antibiotics of last resort: *Capiti-E* and *Capiti-C*, respectively. *Capiti-E* has learned the functions of the previously compiled nefarious protein list (supplemental table 1, (Wittmann et al. 2025)). *Capiti-C* targets were compiled by taking the 2025 AWaRe classification of antibiotics from WHO (“The Selection and Use of Essential Medicines, 2025: WHO AWaRe (Access, Watch, Reserve) Classification of Antibiotics for Evaluation and Monitoring of Use” 2025), subsetting to “antibiotics of last resort”, then intersecting with the CARD database ((McArthur and Wright, n.d.), retrieved 2026-04-24 10:25AM EST) to obtain a final “C” target list of proteins conferring resistance to antibiotics of last resort. It is the largest of the models at 224 targets and demonstrates the scalability of our approach. All three are bundled with the package as ONNX exports.

In addition to the core models, we wrote several helper utilities into *Capiti*. Specifically, capiti-watch bundles several ancillary functions with the model. When invoked from the shell on the raspberry pi in the test rig, it decodes nucleotide bus traces in real time, interprets, and interrupts if a nefarious sequence is detected at high confidence by sending a pulse to the kill wire.

## Results

We find that *Capiti* can reliably distinguish putative pathogenicity from sequence, and outcompetes other methods (Fig 4). (Performance on different subsets is reported in Figure S2.) *Capiti* also works effectively on incompletely synthesized sequences. Unsurprisingly, the true positive rate increases as the model sees more of the sequence. Fig. 5 shows a curve for *Capiti E* demonstrating how true positive rate increases as a function of fraction of protein length seen. There is a noticeable elbow at around 40%, and many targets saturate long before reaching 100%. Intriguingly, there is substantial variability: some proteins can be recognized early, whereas others need much more context before *Capiti* can reliably call them. Figure S3 shows how this kind of information is used: emulated synthesis proceeds until Capiti recognizes a possible nefarious function and ends the run. Figure S4 shows how *Capiti* performs compared to other methods on various prefix lengths.

**Fig 4:**
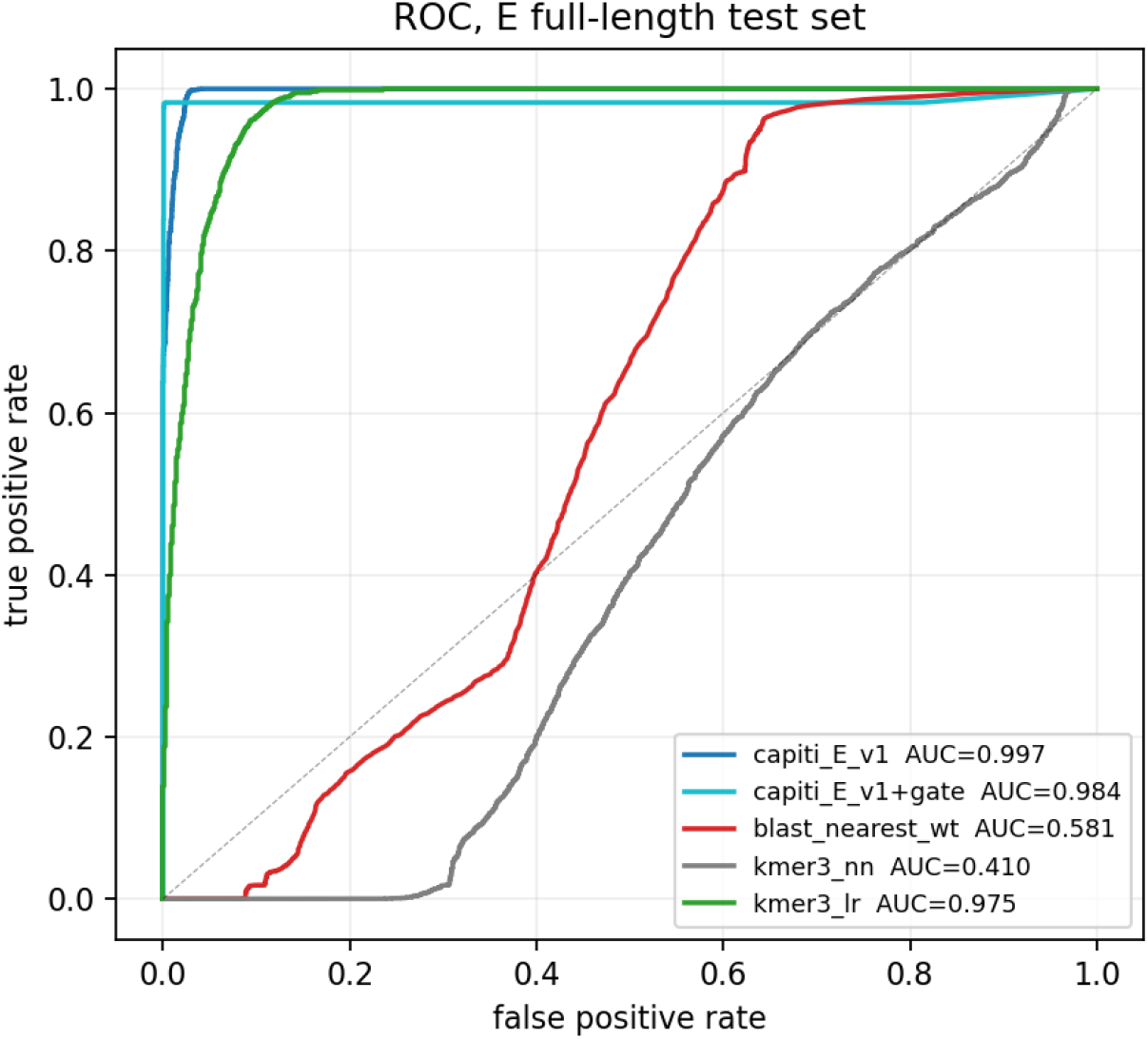
ROC curve of different metrics for identifying E variants. *Capiti* **gate** is a small model modification that can improve performance on `ala_scan`, by forcing probability to zero when active-site mutations are identified by the model. It is togglable via command line flag. blast_nearest_wt is blastp against target sequences. `kmer3_nn` is cosine similarity to 3-mer vectors. `kmer3_lr` is a logistic regression on the same.

**Fig 5:**
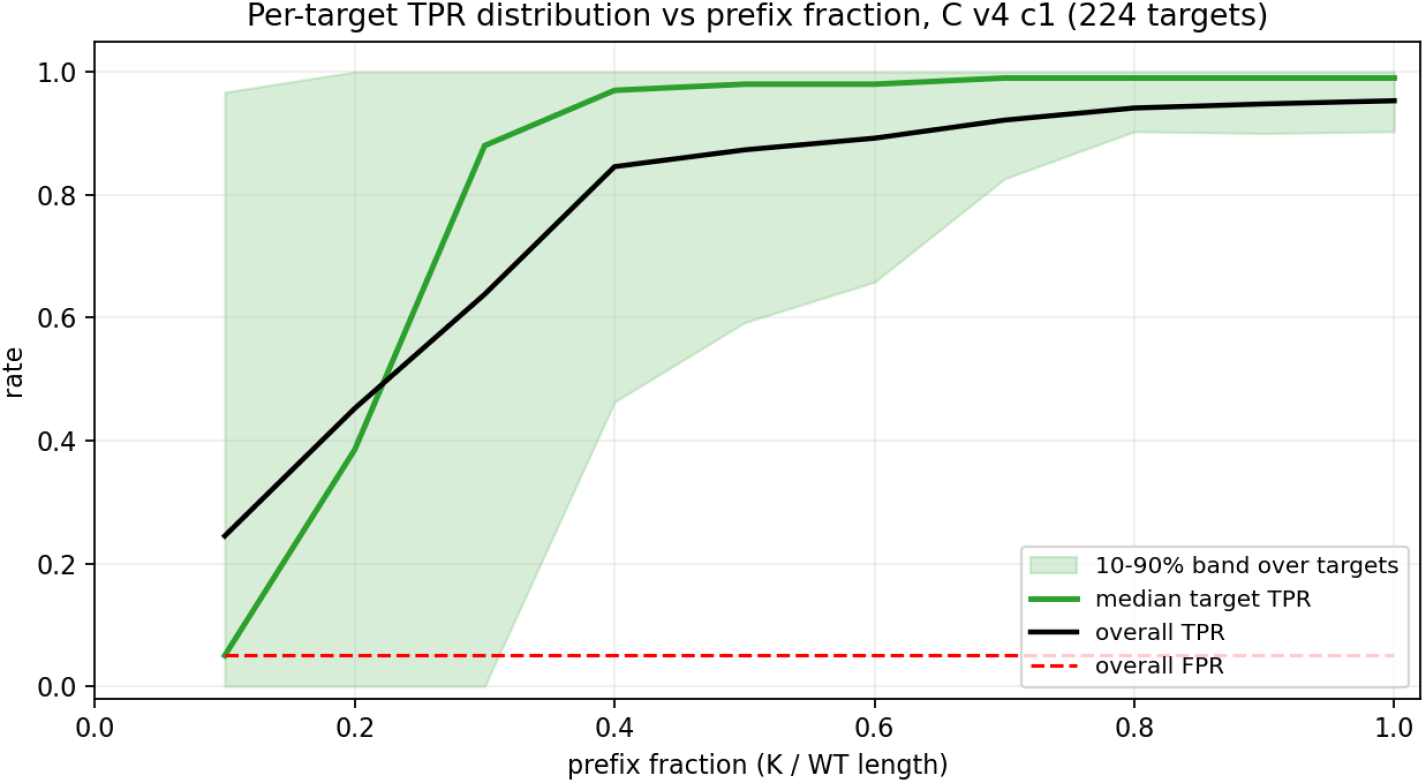
True positive rate as a function of fraction of protein length.

Additionally, we find that *Capiti* can run fast enough to interrupt active synthesis runs, even when emulation is occurring at many fold greater speed than would be possible for a real machine. Click here to see a video of *Capiti* interrupting an emulated synthesis run of a recognized sequence. Click here to see a video of a benign sequence passing through uninterrupted.

### Putatively pathogenic sequences that are not caught by current tools

We ran the *Capiti-E* and *Capiti-C* test sets through two widely used biosecurity screening platforms, SeqScreen and IBBIS commec, and compared them to *Capiti* (Balaji et al. 2022); (Laird et al. 2025a).

Protein sequences were first converted to DNA by substituting each amino acid with its most frequent codon in *Staphylococcus aureus*, using codon frequencies from the Kazusa codon usage database (“Kazusa *Staphylococcus Aureus* Codon Table,” n.d.). *S. aureus* was selected as the reference organism because it is one of the leading sources of antibiotic-resistant bacterial-associated morbidity and mortality worldwide (Taylor et al., 2025).

Note that the underlying positive rate in the E and C datasets were 60.3% and 55.5%, respectively. Table 1) SeqScreen was run on default settings in fast mode on 32 threads. IBBIS commec was run using the light biorisk option. This Capiti model was run without the gate. FNR = False negative rate. FPR = False positive rate.

**Table 1.** Performance of screening software on test dataset (*Capiti* at youden-optimal point).

| Database |  | IBBIS commec | SeqScreen | *Capiti* -gate |
| --- | --- | --- | --- | --- |
| E (known pathogenic proteins) | % functional variants passed (FNR) | 18.47% | 44.69% | 0.24% |
|  | % negative control sequences passed (FPR) | 53.72% | 34.14% | 2.93% |
| C (AMR genes of last resort) | % functional variants passed (FNR) | 95.67% | 40.17% | 0.19% |
|  | % negative control sequences passed (FPR) | 2.14% | 38.03% | 6.60% |

Note that the underlying positive rate in the E and C datasets were 60.3% and 55.5%, respectively. Table 1) SeqScreen was run on default settings in fast mode on 32 threads. IBBIS commec was run

As expected, IBBIS commec failed to flag the majority of *Capiti-C* variants, as AMR genes are likely not included in its threat training set. IBBIS commec passed 18.47% of *Capiti-E* pathogenic variants without flagging them, which raises a significant point of concern. These are sequences created to have the same function as pathogen-associated threats, and a tool designed for biosecurity screening should be expected to catch them.

SeqScreen performed poorly across both datasets, passing 44.69% of *Capiti-E* variants and 40.17% of *Capiti-C* variants. This is particularly notable given that SeqScreen incorporated functional annotations and FunSoC-based pathogenicity predictions. Despite this, nearly half of our putatively pathogenic *Capiti-E* variants evaded detection entirely.

Both tools also showed moderately high false positive rates. Most incorrectly flagged sequences fell into two categories: active site knockouts (ala_scan) and sequence-perturbed variants (perturb30). The false positive rates of both tools have practical implications for researchers. Protein families that share homology with known pathogens (like including antimicrobial peptides, immune components, and conserved enzymes) are likely to be incorrectly flagged despite posing no genuine threat. This places an undue burden on researchers working in these areas, who may face repeated screening rejections for legitimate sequences. If screening tools cannot reliably distinguish benign homologs from functional threats, they risk becoming an obstacle to routine research rather than a meaningful biosecurity safeguard.

### Limitations, Discussion, and corresponding proposed future work

1. The biggest assumption of *Capiti* is that ProteinMPNN created sequences are likely to retain function. A better way to evaluate this *in silico* would be actual structure prediction and comparison to original target structures, producing folding scores. A better version of *Capiti* could be trained on folding scores. We have started work on relatively lightweight validation with ESMfold (Lin et al. 2023), but did not have sufficient GPU hours before the hackathon deadline. (Notably, *in vitro* validation would be technically feasible and theoretically optimal, but carries *severe* risk potential which precludes actual execution.)
2. *Capiti* learns pathogenic function from known pathogenic sequences. This matches the abilities of current open-source tools (and so matches the dominant threat models), but totally novel pathogenic protein design may be possible in the near future. If so, the obvious patch would be to use such tools to create libraries to train better versions of *Capiti*. It *may* be possible to detect “general pathogenic features” (e.g. irreversible binding to essential proteins), in which case the *Capiti* approach would be viable. However, it may be that the space of possible pathogenic sequences is too vast for our simple classification approach. This would require substantial architectural changes. (Theoretically, “truly general pathogenesis prediction” must be possible, since there already exists a system which can predict pathogenicity (living organisms) but whether it is feasible *in silico* is an open question).
3. *Capiti* addresses some specific threat models and not others. Notably, there is currently nothing to prevent a bad actor with physical access to a DNA synthesis machine from simply physically removing *Capiti*. Some hardware changes could make this more difficult (physically welding in the device, running *Capiti* on a dedicated chip embedded in the fluidics microcontroller, etc) and could make assignation of blame after a biosecurity event possible (encrypted tamper-proof logging, breakable plastic seals, etc). Ultimately, however, it is impossible to prevent a sufficiently determined bad actor from simply building their own DNA synthesis machine. It is only possible to make it challenging by preventing dual-use of existing devices, and restriction reagents and equipment (potentially achievable through policy, but outside the scope of the present work).
4. With additional time, *Capiti* could be hardened to several additional attack vectors by applying preprocessing steps at inference. For example, substitution attacks (Adam and McArthur 2024) could be guarded against by running the model on all possible permutations. This would only increase the computational burden and false positive rate by 24-fold, (assuming no swaps *during* a run). Another simple preprocessing at training time could make *Capiti* strand agnostic.
5. *emusynth* emulates the Applied Biosystems 3400 DNA Synthesizer. We felt this was a reasonable first target, but it is hardly the most realistic threat, given its short maximum sequence length. Unfortunately, we had difficulty obtaining technical specifications for more recent, more powerful synthesizers, as they are proprietary.
6. *emusynth* currently only supports one-column synthesis. It could be expanded to multi-column synthesis to more realistically emulate voltage traces, and more realistically challenge capiti-watch.
7. *emusynth* currently uses unrealistic voltages for solenoid signals. We could augment our test-rig with opto-isolators, then monitor 12 or 24 volt lines. This would be a more realistic emulation, and would allow us to deploy *Capiti* directly on a wide variety of real DNA synthesis machines.
8. *Capiti* currently does not have any encrypted logging, a planned feature. The ATECC608 chip did not arrive in time.

## Code and Data

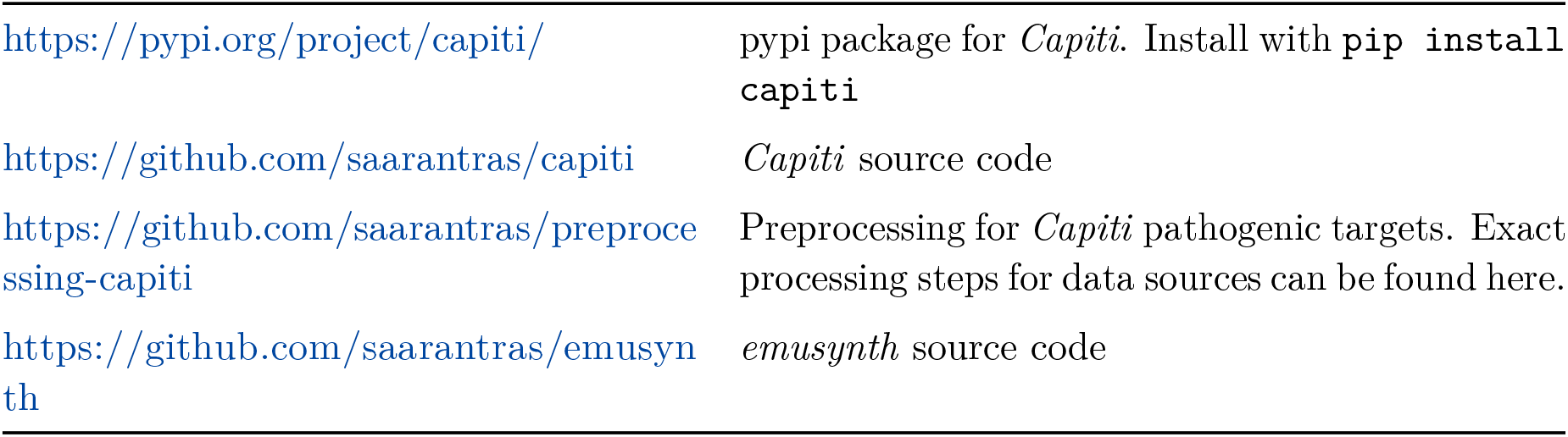

|  |  |
| --- | --- |
| https://pypi.org/project/capiti/ | pypi package for *Capiti*. Install with `pip install capiti` |
| https://github.com/saarantras/capiti | *Capiti* source code |
| https://github.com/saarantras/preprocessing-capiti | Preprocessing for *Capiti* pathogenic targets. Exact processing steps for data sources can be found here. |
| https://github.com/saarantras/emusynth | *emusynth* source code |

Models are bundled with pypi install and ready for inference right out of the box. Generated protein variant libraries possibly available upon request, if they are confirmed not to be dangerous, see appendix.

### Video links

|  |  |
| --- | --- |
| Click here to see a video demonstration of emusynth. | https://www.youtube.com/watch?v=s1grm5U4mPM |
| Click here to see a video of Capiti interrupting an emulated synthesis run of a recognized sequence. | https://www.youtube.com/watch?v=B8rq5Fz7d-c |
| Click here to see a video of a benign sequence passing through uninterrupted. | https://www.youtube.com/watch?v=xCEiOYLIuHU |

## Author Contributions

M.N. conceptualized and led the project, created the presented software & hardware. G.O. performed external tool validation, discovered potential vulnerabilities, and contributed antibiotic resistance expertise. Both authors contributed to writing and reviewed the final manuscript.

## Appendix

### Dual Use Considerations

We expect the dual use potential of *Capiti* and *emusynth* to be minimal to none. Our AI-redesigned pathogenic protein variants which escaped detection by existing tools has the potential for dual use, however this is pending verification with more detailed structural studies, as per the limitations section. Even if the protein variant libraries are hazardous, they can be trivially regenerated from existing open-source software, so we expect that the additional risk from this work is minimal. Still, we do not provide them publicly, erring on the side of safety.

## Supplemental figures

**Fig S1:**
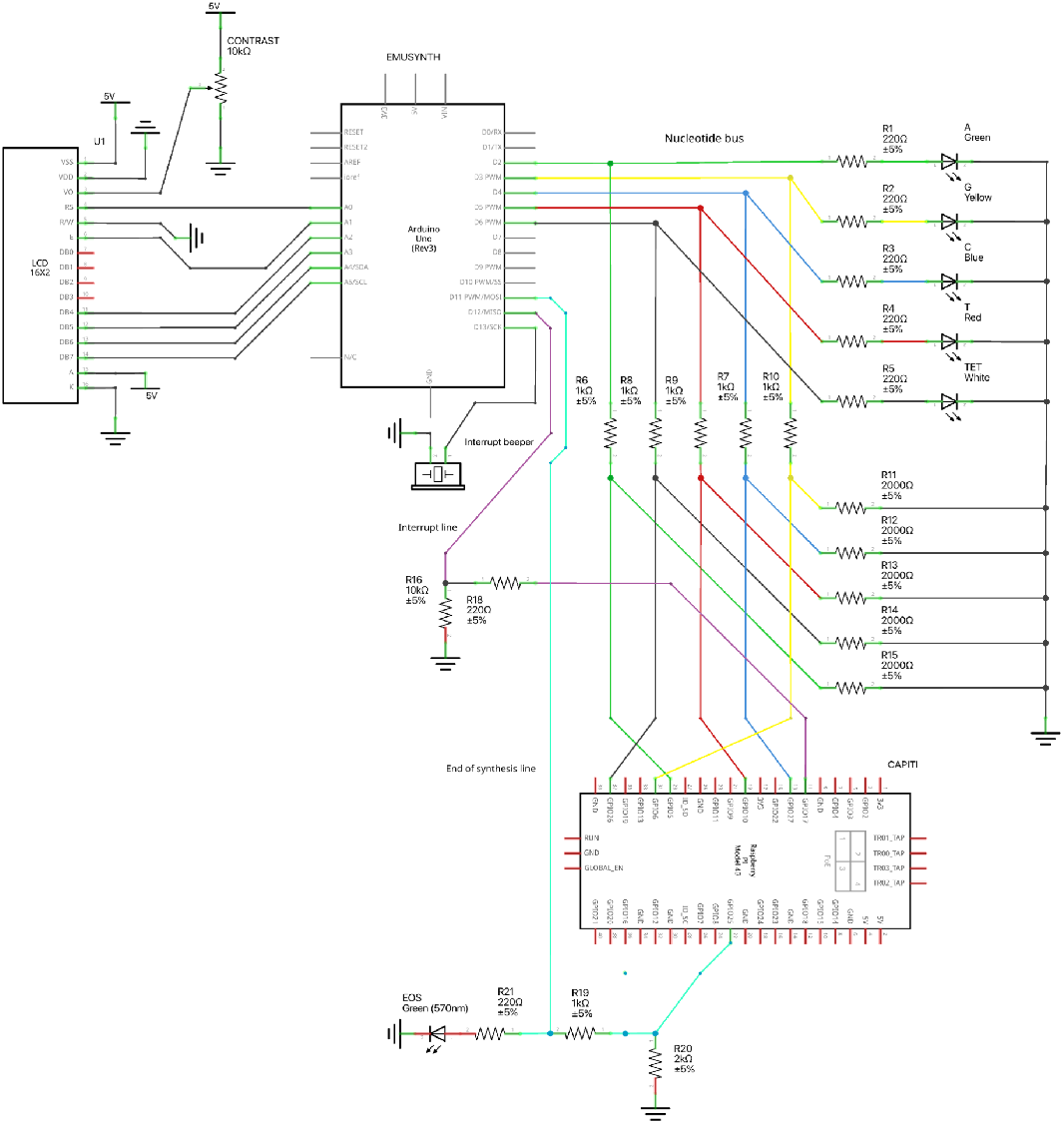
Test rig schematic. Schematic made with Fritzing.

**Fig S2:**
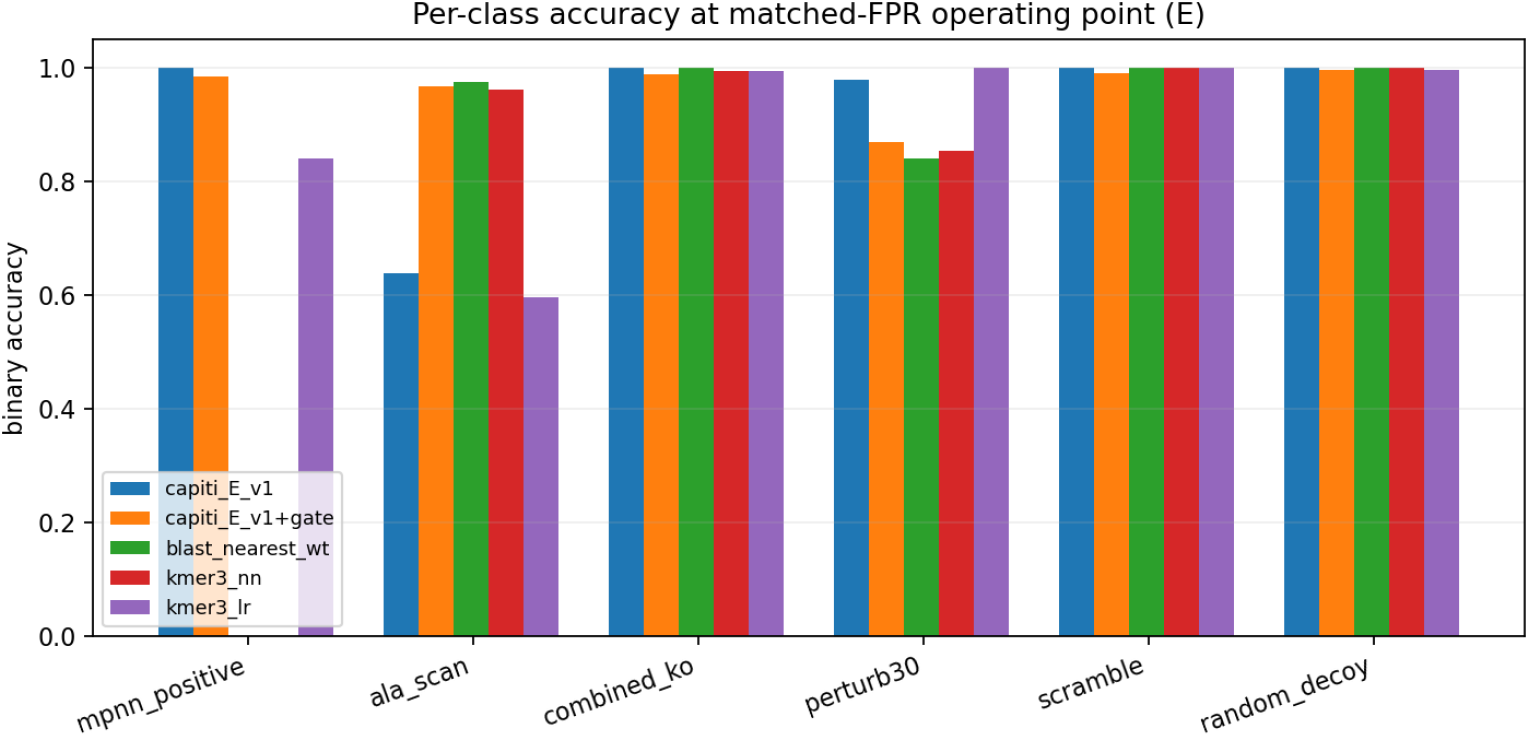
Per-class accuracy for E dataset. Per-class prediction accuracy, calibrated at FPR=5%.

**Fig S3:**
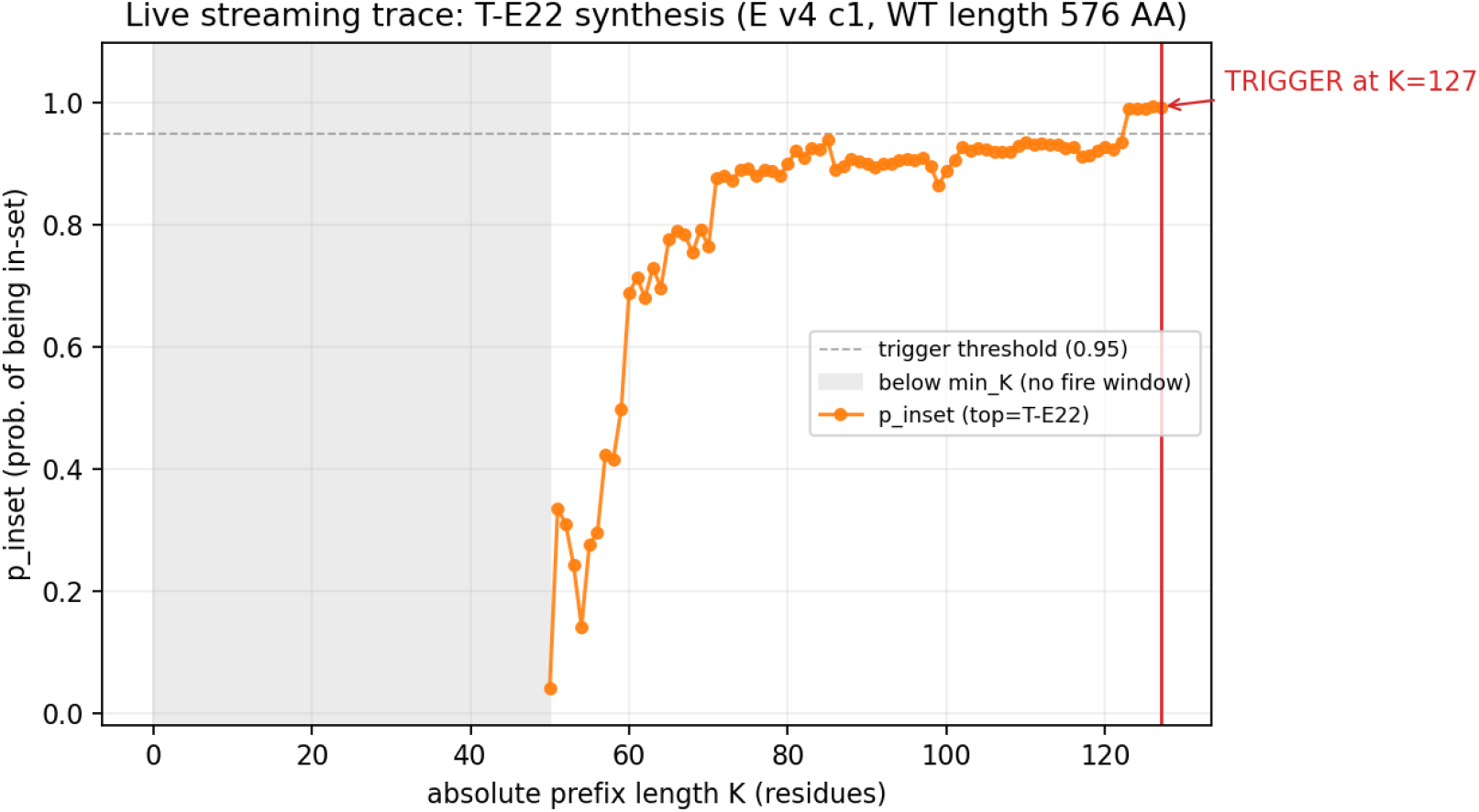
Example trace. Using model *Capiti-E*, target 22. The delay between p_inset exceeding threshold and run kill is caused by a filter: Killing synthesis requires five consecutive residues to exceed threshold, to prevent stochastic drifts above threshold from erroneously killing runs.

## LLM Usage Statement

Claude, under human supervision, assisted with software development.

## Disclaimer

This work was conducted independently by the authors in their individual capacities and was not undertaken as part of their duties or responsibilities at Yale University. The views expressed are those of the authors and do not necessarily reflect those of Yale University.

